# Revelation of a Novel Protein Translocon in Bacterial Plasma Membrane

**DOI:** 10.1101/121335

**Authors:** Feng Jin, Zengyi Chang

## Abstract

Many proteins are translocated across biomembranes via protein translocons in targeting to their subcellular destinations. Hitherto, the SecYEG/Sec61 translocon, existing in prokaryotes and eukaryotes, represents the most intensively studied one. According to the current perception, both periplasmic and β-barrel outer membrane proteins (β-barrel OMPs) are translocated via the SecYEG translocon in bacterial cells, although direct living cell evidences remain lacking. Here, mainly via *in vivo* protein photo-crosslinking analysis, we revealed that the never reported membrane-integrated SecA^N^ protein apparently functions as the translocon for β-barrel OMPs. Additionally, SecA^N^ contains a GXXXG motif known for mediating protein interactions in biomembranes, and processing of β-barrel OMP precursors was severely affected in cells producing an assembly-defective SecA^N^ variant resulted from the GXXXG motif mutations. Furthermore, SecA^N^ was demonstrated to directly interact with the Bam complex, thus likely be a part of the supercomplex that we revealed earlier to be responsible for β-barrel OMP biogenesis.

## Introduction

Proteins are all synthesized by free or membrane-bound ribosomes in the cytosol despite of their diverse final subcellular destinations. One important aspect of protein research has been to decipher how newly synthesized proteins are targeted to their diverse final locations, inside or outside a cell. One early breakthrough in understanding how protein targeting occurs was the realization that these processes are guided by structure information contained within the transported proteins themselves, such as the “signal sequences” (*1-3*), the existence of which was implicated in the “signal hypothesis” initially proposed (*4*). During targeting to their final destinations, many nascent proteins would have to be transported across cellular membranes (*5-8*). In this regard, one early puzzle was how such large and charged structures could traverse the tightly sealed biomembranes without compromising the integrity of the latter. In the early days, it was debated whether the nascent polypeptides travel through the biomembranes in a spontaneous or facilitated manner. Decades of studies revealed that the translocations are escorted by specific “protein-conducting channels” or “translocons” (*9*). Undoubtedly, in studying the biogenesis of proteins, one of the most important but difficult aspects was to identify and characterize the membrane-integrated protein translocons that deliver a specific type of nascent proteins across the hydrophobic membrane barrier. In this regard, what needs to be emphasized is that it remains a great challenge to identify and to learn about the structure, function and mechanism of any membrane-integrated protein in general, especially in living cells.

Among the known protein-conducting channels, the Sec translocon has been the most intensively studied and best understood. It is composed of the core SecYEG heterotrimer in prokaryotic cells or Sec61αβγ heterotrimer in eukaryotic cells, besides other associated proteins (*6, 10*). Briefly, the Sec translocon have been identified and characterized mainly as follows. First, early *in vitro* membrane reconstitution studies hinted the presence of specific protein-conducting channels in mammalian endoplasmic reticulum (ER) membranes (*1, 2*). Second, genetic studies, especially with yeast and Gram-negative bacteria, revealed multiple genes whose mutation affected the protein export process. Out of these, *secA* (*11*) and *secY* (*12, 13*) were identified as key genes whose defect significantly reduced protein secretion in bacteria, and later *sec61α* was characterized as a similar key gene in yeast (*14*). Third, *in vitro* electrophysiological analysis of membrane fragments derived from the mammalian ER (*15*) or bacterial cytoplasmic (*16*) membrane provided further direct evidence supporting the speculation on the existence of protein-conducting channels. Fourth, *in vitro* chemical crosslinking analysis demonstrated direct interactions between protein precursors and the presumed protein translocons, as performed in reconstituted systems (*17*). Fifth, structure determination indicated that the Sec61α subunit in Sec61αβγ and SecY in SecYEG are apparently able to form a central protein-conducting channel (*18-20*). Furthermore, such protein translocation process also consumes energy, either derived from ATP hydrolysis and/or directly from the transmembrane proton gradient (*21*). More recent studies revealed that some proteins were translocated across the cytoplasmic membrane of the bacterial cells apparently via the Tat system, rather than the Sec translocon (*22*).

The envelope of Gram-negative bacteria is composed of two membranes, the cytoplasmic/inner membrane and the outer membrane. Most of the proteins present in the outer membrane belong to the type of β-barrel outer membrane proteins (β-barrel OMPs), because they primarily comprise of β-sheets and adopt a unique cylindrical barrel-like topology (*23, 24*). The β-barrel OMPs, together with the soluble proteins present in the periplasm (the area between the inner and outer membranes) have been considered to be translocated across the inner membrane via the SecYEG translocon (*6, 25, 26*). Meanwhile, it has been recognized that the SecYEG translocon also relies on the cytoplasmic SecA protein, a molecular motor, for providing the driving force. Furthermore, the nascent β-barrel OMPs are known to be further facilitated by more protein factors beyond the inner membrane (*27, 28*). These protein factors include, for example, the periplasm-located primary chaperone SurA (*29-33*) and the outer membrane-located essential β-barrel assembly machine (BAM) complex, in which the BamA protein is a major component (*34-36*). The BamA protein is integrated into the outer membrane via its C-terminal part and extends into the periplasm via its N-terminal POTRA domains (*37, 38*). Our recent studies, mainly via *in vivo* protein photo-crosslinking analyses, unveiled a protein supercomplex that spans the inner and outer membranes and mediates the biogenesis of β-barrel OMPs (*33*).

Besides, β-barrel OMPs are also present in the outer membranes of both mitochondria and chloroplasts in eukaryotic cells. Similarly, they are all synthesized by the cytoplasmic ribosomes and are subsequently targeted to the outer membranes, but are facilitated by the Tom or Toc protein complexes located in the outer membranes (*39, 40*). In this regard, we noticed that, though being a homologue of the bacteria SecYEG (*7, 41-43*), the eukaryotic Sec61αβγ complex in the ER membrane is involved in the biogenesis of proteins dramatically different from the β-barrel OMPs. This in part aroused our interest to learn more about the events of β-barrel OMP biogenesis in living Gram-negative bacterial cells. Again, we mainly performed *in vivo* protein photo-crosslinking studies as mediated by genetically incorporated unnatural amino acid (*44*), a common approach employed in our laboratory (*33, 45-47*).

In this study, we revealed a never reported new protein which seems to function as the translocon (or protein-conducting channel) for nascent β-barrel OMPs. Specifically, we first observed that nascent β-barrel OMPs did not, although nascent periplasmic proteins did, directly interact with SecY, the presumed channel-forming subunit of the SecYEG translocon, in living cells. Later, we accidentally found that the nascent β-barrel OMPs did, but the nascent periplasmic proteins did not, actually interact with a shortened version of SecA that consists of the ~ 45 kD N-terminal region of the full length SecA (of ~ 102 kD), and was thus designated as SecA^N^. We subsequently demonstrated that SecA^N^ specifically existed in the membrane fraction and apparently as homo-oligomers. Meanwhile, along the putative amino acid sequence of SecA^N^, we identified a putative transmembrane domain that contains a GXXXG motif, which has been known to mediate the self-association (i.e., homo-oligomerization) of membrane-integrated proteins. Further, we provided evidences to show that the processing of β-barrel OMP precursors, but not of periplasmic protein precursors, was severely affected in cells producing an assembly-defective SecA^N^ variant due to mutations introduced in the GXXXG motif. Moreover, we demonstrated that the full length form of SecA functions upstream the SecA^N^ or SecYEG translocons. Lastly, we revealed that SecA^N^ directly interacted with BamA of the β-barrel assembly machine (BAM) complex, apparently in the periplasm.

## Results

### Nascent β-barrel outer membrane proteins do not, but nascent periplasmic proteins do directly interact with SecY in living cells

It has been almost universally accepted that both β-barrel OMPs and periplasmic proteins in Gram-negative bacteria are translocated across the inner membrane via the SecYEG translocon during their biogenesis (*6, 26, 48, 49*). In an attempt to verify this, we analyzed whether nascent β-barrel OMPs were able to directly interact with SecY (that largely constitutes the putative protein-conducting channel of the SecYEG translocon) in living cells. For this purpose, we individually introduced the unnatural amino acid pBpa (*p*-benzoyl-L-phenylalanine) (*44-47*) at 21 somehow randomly selected residue positions along the polypeptide of nascent OmpF (a β-barrel OMP) and then assessed whether any of them was able to form photo-crosslinked products with SecY in living cells. In performing the photo-crosslinking analysis, each pBpa variant of OmpF was respectively expressed in the *ompF*-deleted LY928 *E. coli* strain, one whose genome was modified to encode both the orthogonal amino-acyl tRNA synthetase and the orthogonal tRNA that are needed for pBpa incorporation, as we reported earlier (*33*).

To our great surprise, we failed to detect such a photo-crosslinked product band (be of ~ 88.5 kD) between SecY (~ 48.5 kD) and any of these OmpF variants (~ 40 kD), either via blotting analysis by probing SecY (**Fig. 1A**) or via mass spectrometry identification of proteins isolated as photo-crosslinked products of three pBpa variants of OmpF (with pBpa introduced at residue position 26, 65 or 356). It has been observed that the apparent molecular size of the SecY protein on such gel electrophoresis is usually around 36 kD (as shown in **Fig. 1A**), smaller than its actual size, as often be the case for proteins inserted into membranes by using multiple transmembrane α-helices. But potential photo-crosslinked SecY-OmpF product having an apparent molecular size of either ~ 76 kD or 88.5 kD was clearly undetected (**Fig. 1A**). For comparison, we did successfully detect the photo-crosslinked product (with an apparently molecular size of ~ 83 kD) between SecY and the variants of the periplasmic protein SurA (~ 47 kD) when pBpa was placed in the signal peptide region, for instance at residue position 8 or 12 (lanes 2 and 4 in **Fig. S1A**). Taken together, these results strongly suggested that nascent β-barrel OMPs do not, while nascent periplasmic proteins do, directly interact with the SecYEG translocon in living cells. It should be pointed out that we did not detect any photo-crosslinked product when pBpa was introduced at the mature part (e.g. at residue position 110, 125, 140, or 155) of SurA (lanes 8, 9, 10, 11 in **Fig. S1A**). One plausible explanation for these was that the mature part of SurA interacted with SecY only transiently and thus failed to be captured by the photo-crosslinking technique employed by us.

**Figure 1.**
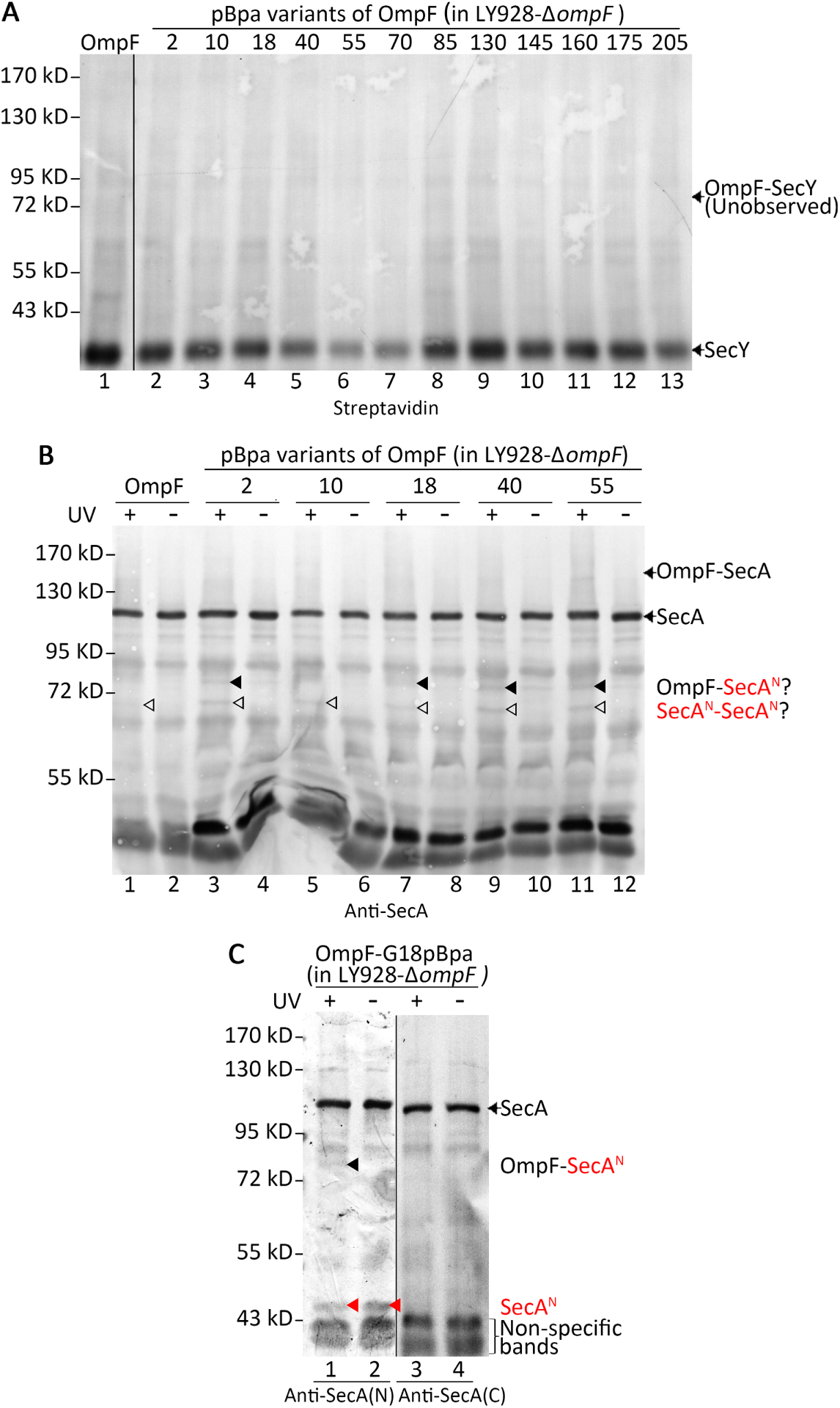
Nascent β-barrel OMPs do not interact with SecY, but with a shortened version of SecA (SecA^N^). (**A, B**) Blotting results for the detection of the *in vivo* protein photo-crosslinked products of the indicated pBpa variants of OmpF that were expressed in the LY928-Δ*ompF* strain, probed with streptavidin AP-conjugate against the Avi-tag linked to SecY (**A**), or with antibodies against the full length SecA (i.e., a mixture of antibodies against the N-terminal fragment 1-209 and those against the C-terminal fragment 665-820 of SecA) (**B**). (**C**) Immunoblotting results for the detection of the photo-crosslinked product between OmpF-G18pBpa and SecA^N^, as well as free SecA^N^, probed with antibodies against the N- (lane 1) or C-terminal (lane 3) region of SecA. All protein samples were resolved by SDS-PAGE before subjecting to the blotting analysis. Residue positions for OmpF are numbered by including the signal peptides. Samples of cells expressing wild type OmpF (with no pBpa incorporation) were analyzed as negative controls (lane 1 in **A**, and lanes 1 and 2 in **B**). Indicated on the right of each gel are positions of the indicated protein monomers or photo-crosslinked products, and on the left are positions of the molecular weight markers.

In agreement with these *in vivo* protein photo-crosslinking results, our genetic studies also revealed that the processing of precursors of β-barrel OMPs (as represented by OmpF and OmpA) was far less defective (i.e., with far less precursors accumulated) compared with that of periplasmic proteins (as represented by SurA and the maltose binding protein, MBP) in the cold sensitive *secY39* mutant cells (*50*) that was cultured at the non-permissive temperature of 20°C and thus possessing a defective SecY protein (**Fig. S2A**). This indicated that, in physiological terms, the processing of precursors for β-barrel OMPs relied on the SecYEG transcolon in a far less degree than that for periplasmic proteins. It is worth noting that our results were largely consistent with what was reported before (*50*), where it was observed that this *secY39* mutation affected MBP export more strongly than it affected the OmpA export. Likewise, in another *secY* mutant, *secY125*, the translocation of periplasmic proteins MBP and DsbA were also found to be more severely affected than that of the β-barrel outer membrane protein OmpA (*51*). Collectively, these data suggested that the SecY channel may translocate the nascent periplasmic proteins, but unlikely the nascent β-barrel OMPs, across the inner membrane in living bacterial cells.

### Nascent β-barrel outer membrane proteins do, but nascent periplasmic proteins do not directly interact with a shortened version of SecA (SecA^N^) in living cells

For comparison and as a kind of positive control, we also assessed whether SecA, which has been demonstrated to play a role in translocating both nascent β-barrel OMPs and nascent periplasmic proteins by its presence in the cytosol or as associated with the inner membrane (*52*), could be photo-crosslinked with these two types of proteins in living cells. To achieve this goal, we made use of the same pBpa variants of OmpF and SurA that were examined above, and probed with antibodies against SecA. The immunoblotting results shown in **Figs. 1B** and **S1B** revealed the following. First, we hardly detected any photo-crosslinked product between SecA and OmpF or SurA at the expected size position, except seeing a rather faint band of ~ 142 kD for the OmpF variant in which pBpa was introduced at residue position 55 (lane 11 in **Fig. 1B**). Second, unexpectedly, a photo-crosslinked product band of ~ 85 kD was detected for 4 of the 5 represented pBpa variants of OmpF (lanes 3, 7, 9. 11 in **Fig. 1B**) but undetectable for the wild type OmpF with no pBpa incorporation (lane 1). Third, also unexpectedly, a photo-crosslinked product band of ~ 65 kD was detected for all the UV-exposed cell samples, including those only expressing the wild-type OmpF or SurA (lane 1 in **Fig. 1B** and lane 17 in **S1B**, respectively). At this point, the most plausible explanations on these two photo-crosslinked products are such that the ~ 85 kD band represented a photo-crosslinked product between the pBpa variants of nascent OmpF (~ 40 kD) and a shortened version (presumably of ~ 45 kD) of SecA, in turn the ~ 65 kD band represented a photo-crosslinked dimer of this shortened version of SecA. These preliminary explanations will be further substantiated by multiple other observations described below.

We next attempted to verify whether this presumed shortened version of SecA indeed existed. To achieve this goal, we decided to employ a type of gel electrophoresis that would resolve smaller proteins with a higher resolution (*53*) as well as to probe with antibodies against either the N-terminal or C-terminal region of SecA. To our great interest, we did find that the ~ 85 kD photo-crosslinked product of the OmpF-G18pBpa variant was detected only with antibodies against the N-terminal region (fragment 1-209), but not with those against the C-terminal region (fragment 665-820) of SecA (lane 1 vs 3 in **Fig 1C**). Furthermore, this immunoblotting analysis also clearly unveiled one shortened version of SecA with an apparent molecular mass of ~ 45 kD, which again could only be detected with antibodies against the N-terminal region of SecA (lane 1 vs lane 3 in **Fig 1C**). Of note, we failed to detect photo-crosslinked product between SecA^N^ and any of the 10 pBpa variants of SurA (**Fig. S1B**). Given that this shortened version of SecA is apparently made of the N-terminal region of the full length form of SecA, we hereafter designate it as SecA^N^.

Taken together, these results strongly suggested the existence of the hitherto unreported SecA^N^, which interacted with nascent β-barrel OMPs, but not with nascent periplasmic proteins, in living cells. Additionally, this SecA^N^ seemed to be able to form a photo-crosslinked dimer upon UV irradiation, thus likely existed as a homo-oligomer in living cells. Such a photo-crosslinked dimer of SecA^N^ was plausibly formed through an interacting pair of tyrosine residues at the interface of the interacting monomers, as previously observed in other proteins (*54*). The observation that the apparent size of the photo-crosslinked SecA^N^ dimer was ~ 65 kD, rather than the expected ~ 90 kD (2 X 45 kD), might be due to the fact that the photo-crosslinked product was formed via two interacting tyrosine residues located in the middle of the SecA^N^ polypeptide chain (thus to form a “H”-shaped crosslinked product), as reported before for other proteins (*55*).

### SecA^N^ is detected wholly in the membrane fraction and apparently exists as homo-oligomers

We subsequently attempted to elucidate whether SecA^N^ could indeed function as the protein-conducting channel for β-barrel OMPs. Towards this goal, we first investigated whether SecA^N^ is located in the membrane fraction, a prerequisite for it to function as a transmembrane protein-conducting channel. In particular, we separated the membrane and soluble fractions of the cell lysates by differential centrifugation (*56*) before subjecting each fraction to blotting analysis.

Our immunoblotting results, shown in **Fig. 2**, clearly demonstrated that the SecA^N^ protein was detected almost wholly in the membrane fraction (lanes 1 and 2), hardly any in the soluble fraction (lanes 5 and 6). Interestingly, similar to the membrane-integrated trimeric OmpF (e.g., lanes 2-4 in **Fig. S2A**), SecA^N^ in the membrane fraction was also detected as a smear on the gel when the sample was unboiled (lanes 3 and 4 in **Fig. 2A**), mobilizing with a rate significantly lower than that of its monomeric form, as detected in the boiled samples (lanes 1 and 2 in **Fig. 2A**). This strongly suggested that SecA^N^, like OmpF, existed as a membrane-integrated protein homo-oligomer. This conclusion was further supported by our observations that crosslinked SecA^N^ dimers could also be effectively produced by directly exposing the membrane fraction to UV irradiation (lane 1 in **Fig. 2A**) and that the SecA^N^ protein almost all remained in the membrane fraction after being treated with 8 M urea (lane 3 in **Fig. S3**), that would effectively remove proteins associated to the surface of but not those integrated into the biomembrane (*57*).

**Figure 2.**
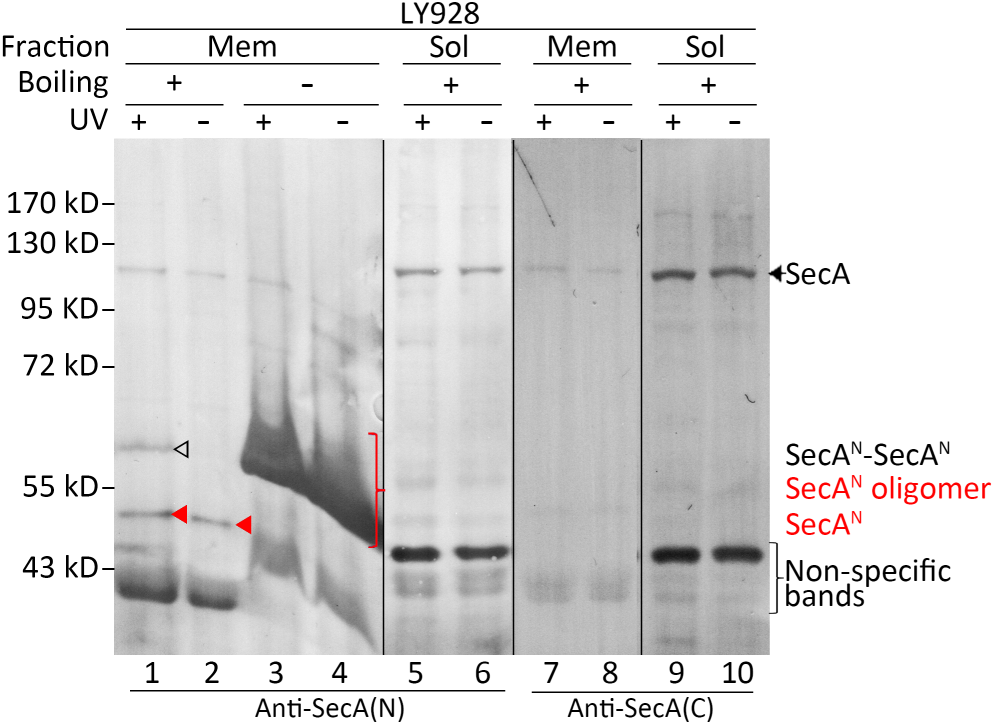
SecA^N^ is detected wholly in the membrane fraction and apparently exists as homo-oligomers. Immunoblotting results for the detection of SecA^N^ in the membrane (Mem) and soluble (Sol)fractions of the bacterial cell lysates, as well as of crosslinked SecA^N^ dimers after exposing membrane fraction to UV irradiation, probed with antibodies against the N- (lanes 1-6) or C-terminal (lanes 7-10) region of SecA. The polymerized separating gel was 10%. Indicated on the right of the gel are positions of SecA, monomeric SecA^N^, SecA^N^ oligomers, photo-crosslinked products of SecA^N^-SecA^N^ dimer, and onthe left are positions of the molecular weight markers.

### SecA^N^ contains a putative transmembrane domain in which a GXXXG motif known for mediating protein-protein interactions in biomembranes is identified

In an attempt to provide more evidences to support our conclusion that SecA^N^ is a membrane-integrated homo-oligomeric protein, we also examined whether there are any putative transmembrane domains along the polypeptide chain of SecA^N^. And further, whether any GXXXG motif, that has been known to mediate self-associations of membrane-integrated proteins (*58, 59*), could be identified in such putative transmembrane domains. In this regard, based on its apparent molecular weight of ~ 45 kD as revealed above (**Figs. 1C** and **2**), we took SecA^N^ as a protein that consists of ~ 420 amino acid residues from the N-terminus of SecA. For these purposes, we subjected the N-terminal 416 residues (~ 47 kD) of SecA to the TMpred software prediction (*60, 61*), as provided by the online server (http://www.ch.embnet.org/software/TMPRED_form.html). As a result, one putative transmembrane domain, being ^145^PLFEFLGLTVGINLPGMPAPA^165^, were clearly identified from the putative SecA^N^ sequence.

Remarkably, we did identify one single GXXXG (being ^151^GLTVG^155^) motif across the whole polypeptide chain of *E. coli* full length SecA, and this motif was indeed present in the predicted transmembrane domain in SecA^N^. Of note, we further revealed that such a GXXXG motif, together the predicted putative transmembrane domain, was also identified in SecA proteins of other bacterial species (**Table 1**). We also noticed that this GXXXG motif (colored red) was located away from the dimerization interface identified in the full length SecA crystal structures, and partially in a β-strand and partially in a loop, as shown in **Fig. S5**. It is worth noting that the structure of SecA^N^, presumably existing as a membrane-integrated homo-oligomer according to our experimental observations described above, would be significantly different from that of the SecA^N^ equivalent in the reported structure of the full length SecA (*62*).

**Table 1.** The GXXXG motif that was identified in a putative transmembrane domain of *E. coli* SecA is highly conserved in other bacterial species.

| Bacterial species | Identity (%)<sup>1</sup> | TM<sup>2</sup> | Score<sup>3</sup> | The motif |
| --- | --- | --- | --- | --- |
| *Escherichia coli* | 100 | 145-165 (21) | 735 | <sup>151</sup>GLTVG<sup>155</sup> |
| *Lactobacillus delbrueckii* | 48 | 143-161 (19) | 443 | <sup>149</sup>GLTVG<sup>153</sup> |
| *Stenotrophomonas maltophilia* | 62 | 141-161 (21) | 722 | <sup>151</sup>GLSVG<sup>155</sup> |
| *Bacillus subtilis* | 50 | 141-159 (19) | 172 | <sup>149</sup>GLTVG<sup>153</sup> |
| *Pseudomonas aeruginosa* | 64 | 145-162 (18) | 310 | <sup>151</sup>GLSVG<sup>155</sup> |
| *Mycobacterium tuberculosis* | 35 | 145-165 (17) | 555 | <sup>158</sup>GLTVG<sup>162</sup> |
| *Streptococcus pyogenes* | 44 | 143-161 (19) | 581 | <sup>149</sup>GLSVG<sup>153</sup> |

<sup>1</sup>Listed here is the percentage of amino acid sequence identity for the SecA of the indicated bacterial species when compared with that of *E. coli* SecA.
<sup>2</sup>TM: Transmembrane region, shown are the residue ranges and the total number of residues (in the brackets) of the predicted transmembrane domain.
<sup>3</sup>Here, we also included the transmembrane domains predicted with a score <500.

### Processing of precursors of β-barrel OMPs is severely affected in cells producing an assembly-defective SecA^N^ variant due to mutations introduced in the GXXXG motif

In an attempt to provide more direct evidences to support our conclusion that SecA^N^ acts as the protein-conducting channel for translocating β-barrel OMPs, we next tried to conduct functional studies, i.e., trying to observe the biological consequences after making SecA^N^ defective but keeping SecA fully functional. We initially reasoned that if SecA^N^ is generated from the full length SecA via the cleavage of a certain protease, we might be able to eliminate SecA^N^ but to keep the full length SecA unaffected in *E. coli* strains lacking a particular protease. We then examined multiple knockout strains (obtained from the Keio Collection, Japan), but found the level of SecA^N^ protein in cells lacking GlpG (a protease integrated in the inner membrane), or HslU, HslV, YcbZ and YifB (proteases present in the cytoplasm) is largely comparable with that in wild-type cells. We then decided to try to generate SecA variants by replacing some of the residues in the GXXXG motif (^151^GLTVG^155^), hoping that some of them would generate assembly-defective (thus nonfunctional) SecA^N^, while keeping the full length SecA fully (or at least largely) functional.

Given that SecA is essential for cell growth, we would have to evaluate the structure and function of these SecA^N^ variants in a temperature sensitive mutant strain under the non-permissive temperature when its endogenous SecA and SecA^N^ become defective. In this regard, we constructed the LY928-*secA81* strain, in which the genomic *secA* gene was replaced by the *secA81* gene that encodes a temperature sensitive SecA-G516D (or SecA81) mutant protein, being functional at the permissive temperature of 37°C but defective at the non-permissive temperature of 42°C (*63*). We then revealed that, when the LY928-*secA81* cells were first grown at the permissive temperature of 37°C for 6 h to an OD_600_ of 0.8~1.0, and then shifted to and further incubated at the non-permissive temperature of 42°C for as short as 1 h, the SecA^N^ protein present in the cells remained largely as effectively assembled homo-oligomer forms, thus remained functional (**Fig. S4A**). However, when the LY928-*secA81* cells were first grown at the permissive temperature of 37°C for 3 h to an OD_600_ of 0.3~0.4, and then shifted to and further incubated at the non-permissive temperature of 42°C for as long as 3 h, the SecA^N^ present in the cells were largely unassembled monomers (i.e, being detectable in unboiled cell samples), thus being structurally and functional defective (**Fig. S4B**). These observations indicated that the LY928-*secA81* cells that were incubated at the non-permissive temperature for about 3 hours could be applied for evaluating the structural and functional state of the SecA^N^ variants that we planned to generate. These results meanwhile implicated that the formation of a functional SecA^N^, although the detail mechanism of which remained unknown, apparently relied on the presence of a well folded full length SecA.

Furthermore, in order to distinguish the SecA^N^ variants from the endogenous SecA^N^, we tried to add an Avi-tag onto the SecA^N^ protein before we generated the variants. For this purpose, we initially tried to add the tag to the N-terminus of SecA, but the fused SecA protein failed to be produced in cells. We then tried to insert the Avi-tag into a loop region between residues Ala229 and Glu230 (*62*), and successfully produced the fusion protein that we designated as Avi-SecA. We further demonstrated that this Avi-SecA protein was fully capable in rescuing the growth (marked as WT in **Fig. S6**), in producing the assembled SecA^N^ oligomers (lanes 6 in **Fig. 3A** and lanes 13 in **Fig. 3B**), and in facilitating the processing of β-barrel OMP precursors (lane 9 in **Fig. 3C**) for the LY928-*secA81* cells cultured at the non-permissive temperature of 42°C.

**Figure 3.**
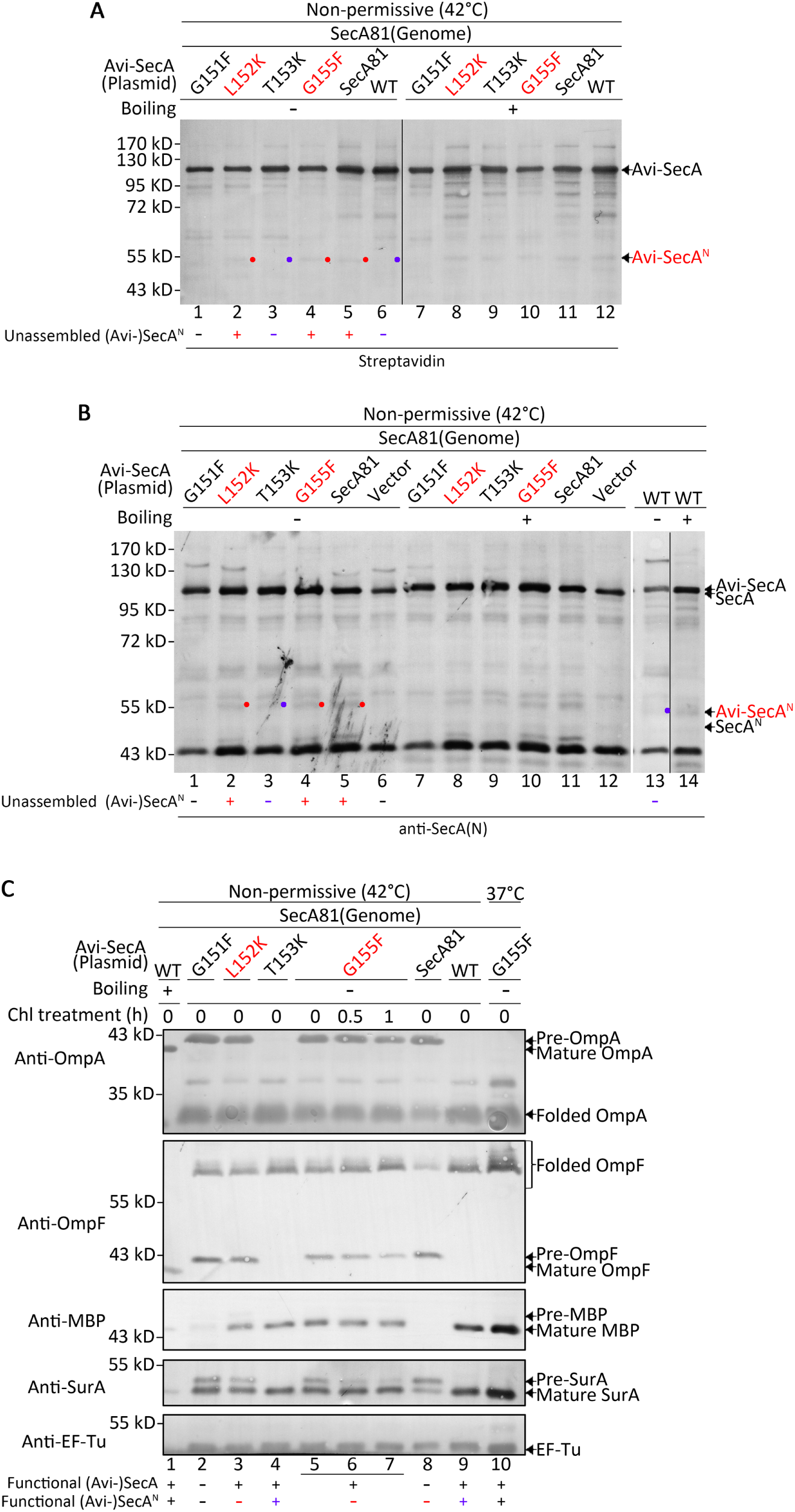
Processing of precursors of β-barrel OMPs, but not of periplasmic proteins, was severely affected in cells failing to produce a functional oligomeric SecA^N^ due to a mutation in the GXXXG motif. (**A, B**) Blotting results for the detection of Avi-SecA and Avi-SecA^N^ monomers in LY928-*secA81* cells transformed with plasmids expressing the indicated variants of Avi-SecA, probed either with streptavidin-AP against the Avi-tag (**A**) or antibodies against the N-terminal region of SecA (**B**). (**C**) Immunoblotting results for the detection of the precursor, mature and folded forms of the indicated β-barrel OMPs (OmpA and OmpF) and periplasmic proteins (SurA and MBP). The LY928-*secA81* cells, initially incubated at 37°C to an OD_600_ of ~ 0.3-0.4, were then either incubated at the non-permissive temperature of 42°C (lanes 1-12 in **A**; lanes 1-14 in **B**; lanes 1-9 in **C**), or continuously at the permissive temperature of 37°C (lane 10 in **C**), both for 3 hours; the cells were treated with chloramphenicol (Chl) for the indicated duration in performing the chasing experiments (lanes 5-7 in **C**). Protein samples from LY928-*secA81* cells transformed with a plasmid either expressing the mutant SecA81 (lanes 5 and 11 in **A** and **B**, and lane 8 in **C**) or the wild type SecA protein (lanes 6 and 12 in **A**, lanes 13 and 14 in **B**, and lane 9 in **C**) were analyzed as negative and positive controls, respectively. Boiled protein samples of wild type cells (WT) were analyzed for indicating the positions of the mature forms (which lacked the signal peptide and thus mobilized with a slightly higher rate than the precursors) of β-barrel OMPs and periplasmic proteins (lane 1 in **C**). The desired mutations (i.e., those that generated a normal full length Avi-SecA but a defective Avi-SecA^N^) were colored red. The red dots indicate cases where unassembled Avi-SecA^N^ monomers were detected in the unboiled samples, while the blue dots indicate cases where unassembled SecA^N^ monomer was undetected in the unboiled samples (in panels **A** and **B**). The EF-Tu protein was analyzed here to indicate an equal loading of cell lysate samples in lanes 2-10 in **C**. The polymerized separating gel was 10%. Indicated on the right of the gel are the positions of different forms of β-barrel OMPs (precursor, mature and folded), periplasmic proteins (precursor and mature), Avi-SecA, and Avi-SecA^N^ monomer, and on the left are positions of the molecular weight markers. Protein samples were unboiled in performing the semi-native SDS-PAGE analysis (lanes 1-6 in **A**, lanes 1-6 and 13 in **B**, and lanes 2-10 in **C**). Note: we also generated the Avi-SecA-V154K variant, but it was undetectable in the cells and was thus not analyzed here.

We then evaluated the Avi-SecA variants of the GXXXG motif, as expressed from a low copy number plasmid pYLC with the control of *secA*’s natural promoter and SecM (*64*), in the LY928-*secA81* cells. Specifically, the transformed cells were first grown at the permissive temperature of 37°C for 3 h to an OD_600_ of 0.3~0.4, and then shifted to and further incubated at the non-permissive temperature of 42°C for 3 h (**Fig. S4B**) before the properties of the variants were examined. The observed properties of the five GXXXG motif variants of SecA are summarized in **Table 2**) and briefly described below. Firstly, to our great satisfaction, we obtained two variants, Avi-SecA-L152K and Avi-SecA-G155F (colored red in **Table 2**), that failed to produce the assembled Avi-SecA^N^ (i.e., Avi-SecA^N^ monomer was effectively detected in unboiled samples (lanes 2 and 4 in **Figs. 3A and 3B**, marked by red dots) but partially rescued the cell growth (Labeled as L152K and G155F in **Fig. S6**). Remarkably, these two variants rendered the processing of β-barrel OMP (again, represented by OmpA and OmpF) precursors to be significantly retarded such that their precursors were dramatically accumulated, but the processing of periplasmic protein (again, represented by MBP and SurA) precursors was far less affected such that the nascent proteins of SurA and MBP were mostly and largely processed to the mature forms, respectively (lanes 3 and 5 vs 8 in **Fig. 3C**). Moreover, our pulse-chase experiments indicated that after adding chloramphenicol to inhibit protein synthesis in cells expressing Avi-SecA-G155F, the high level of the accumulated OmpF or OmpA precursors were hardly processed while the low level of the accumulated SurA precursor was efficiently processed during the 1 h chasing time (lanes 5-7, **Fig. 3C**). Collectively, these genetic and biochemical data strongly supported our conclusion that the SecA^N^ homo-oligomer most likely functions as the translocon for β-barrel OMP, but not for periplasmic proteins. It should be pointed out that the full length form Avi-SecA produced from these two variants were apparently functional, given that the processing of the periplasmic protein precursors was hardly affected.

**Table 2.**
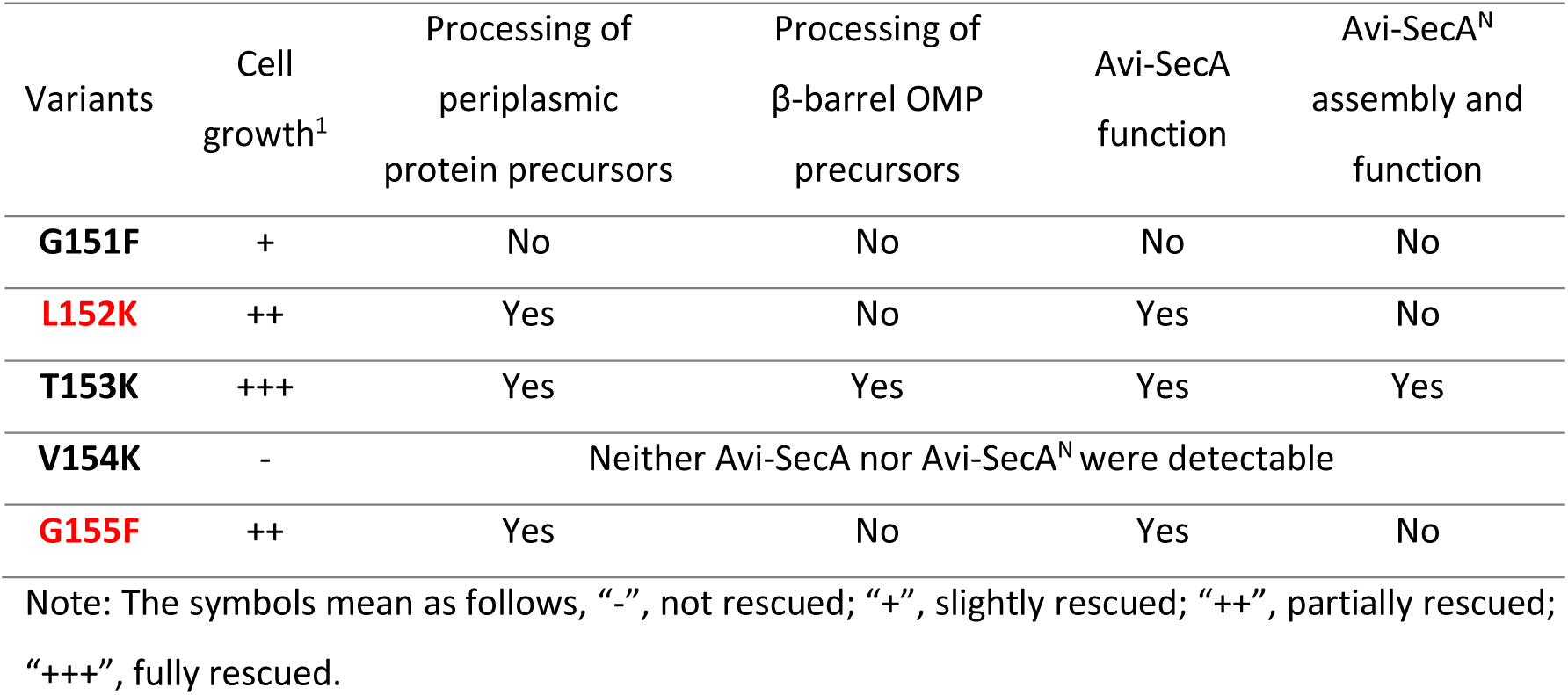
A summary of the functional and structural consequences for replacing the residues in the GXXXG motif of Avi-SecA (based on results displayed in **Fig. 3**).

| Variants | Cell growth<sup>1</sup> | Processing of periplasmic protein precursors | Processing of β-barrel OMP precursors | Avi-SecA function | Avi-SecA<sup>N</sup> assembly and function |
| --- | --- | --- | --- | --- | --- |
| **G151F** | + | No | No | No | No |
| **L152K** | ++ | Yes | No | Yes | No |
| **T153K** | +++ | Yes | Yes | Yes | Yes |
| **V154K** | - | Neither Avi-SecA nor Avi-SecA<sup>N</sup> were detectable |  |  |  |
| **G155F** | ++ | Yes | No | Yes | No |

Note: The symbols mean as follows, “-”, not rescued; “+”, slightly rescued; “++”, partially rescued; “+++”, fully rescued.

Secondly, the Avi-SecA-T153K variant seemed to have produced functional SecA^N^ (and SecA). Specifically, it rescued the cell growth (labeled as T153K in **Fig. S6**), it produced the effectively assembled Avi-SecA^N^ (lanes 3 in **Figs. 3A and 3B**, monomers were undetectable in unboiled samples, as marked by the blue dot), and it allowed the precursors of β-barrel OMPs to be effectively processed (lane 4 in **Fig. 3C**), all being similar to the effects of the wild type Avi-SecA (labeled as WT in **Fig. S6**; lane 6 in **Fig. 3A**; lane 13 in **Fig. 3B**; lane 9 in **Fig. 3C**). Thirdly, the Avi-SecA-G151F variant apparently produced a misfolded Avi-SecA^N^ (which was degraded by the quality control system), as well as a non-functional Avi-SecA. Specifically, it slightly rescued the cell growth (labeled as G151F in **Fig. S6**), it neither produced detectable Avi-SecA^N^ (i.e., its monomeric form was undetectable even in the boiled samples; lanes 1 and 7 in **Figs. 3A and 3B**), and it allowed the precursors of neither β-barrel OMPs nor periplasmic proteins to be effectively processed (lane 2 in **Fig. 3C**). Of note, in case of the Avi-SecA-G151F variant, a lack of functional full length form of Avi-SecA led to the degradation of the precursors of MBP (thus the mature form of MBP became neither detectable, as shown in lane 2 in **Fig. 3C**). Fourth, for the Avi-SecA-V154K variant, we detected the presence of neither Avi-SecA nor Avi-SecA^N^ in the cells, and correspondingly, it did not rescue the growth of LY928-*secA81* (labeled as V154K in **Fig S6**), suggesting that this replacement caused a dramatic structural disturbance on both SecA and SecA^N^, which resulted in their degradation in the cells by the quality control system.

Collectively, these genetic and biochemical data, presented in **Figs. S6**, **3A, 3B and 3C**, demonstrated that the GXXXG motif is indeed important for SecA^N^ to form a functional homo-oligomer, and also strongly suggested that SecA^N^ functions as the translocon for β-barrel OMPs.

### Prolonged upstream interactions of nascent β-barrel OMPs or periplasmic proteins with a defective SecA prevent their downstream interactions with SecA^N^ or SecY, respectively

Our observations described above strongly suggested that the SecA^N^ homo-oligomer functions as the transmembrane conducting channel for nascent β-barrel OMPs, similar to what SecYEG functions for nascent periplasmic proteins. Next we intended to seek for more supporting evidences by investigating the functional relationship between SecA^N^ and other key protein factors involved in the biogenesis of β-barrel OMPs.

Given that the full length SecA has been known to act upstream (i.e., by functioning in the cytoplasm) the SecYEG translocon, we attempted to elucidate whether SecA would also act upstream the membrane-integrated SecA^N^ translocon. Nevertheless, likely due to the transient nature of the interaction between SecA and its client proteins, we failed to detect significant interaction between SecA and the nascent β-barrel OMPs or periplasmic proteins in living cells (as shown in **Figs. 1B** and **S1B**). We decided to repeat these photo-crosslinking experiments with the LY928-*secA81* strain cultured at the non-permissive temperature, hoping that the interaction between the defective SecA81 and its client proteins would be prolonged, which might in turn prevent the occurrence of interactions between these client proteins and the downstream SecY or SecA^N^ translocon. In particular, we expressed the same set of pBpa variants of OmpF and SurA (as analyzed in **Figs. 1B** and **S1B**) in the LY928-*secA81* cells, and first inoculated the cells at the permissive temperature of 37°C to the mid-log phase, and then shifted to and further incubated at the non-permissive-temperature of 42°C for 1 hour before exposing the cells to UV irradiation.

The immunoblotting results revealed a significant formation of photo-crosslinked products between SecA and nascent OmpF, when pBpa was placed at residue positions 10 and 18 (both in the signal peptide) or 55 (in the mature protein) in OmpF (lanes 3, 4 and 6 in **Fig. 4A**), or between SecA and nascent SurA, when pBpa was placed at residue position 8 in SurA (in the signal peptide) (lane 2 in **Fig. 4B**). Of note, we detected two photo-crosslinked product bands between SecA and the nascent OmpF (**Fig. 4A**), with the smaller-sized band to be apparently identical as the one detected in **Fig. 1B** (lane 11). These observations apparently suggested that each nascent polypeptide of OmpF interacted with two different sites in SecA. Whether these two different manners of interactions reflected those between nascent β-barrel OMPs and two forms of SecA (such as the cytoplasmic and the membrane-associated/integrated forms) is undoubtedly worth further investigations. Furthermore, it should be pointed out that we failed to detect any photo-crosslinked product when pBpa was placed in the mature part of SurA (lanes 8, 9, 10, 11, **Fig. 4B**), apparently indicating that SecA interacted with the mature part of periplasmic proteins in a highly transient fashion. It is worth noting that under this non-permissive condition, precursors of neither β-barrel OMPs nor periplasmic proteins were effectively processed in LY928-*secA81* cells, as shown by data displayed in **Fig. S8**, which was consistent with what was reported before (*63*) and in agreement with our photo-crosslinking results shown here (**Fig. 4**).

**Figure 4.**
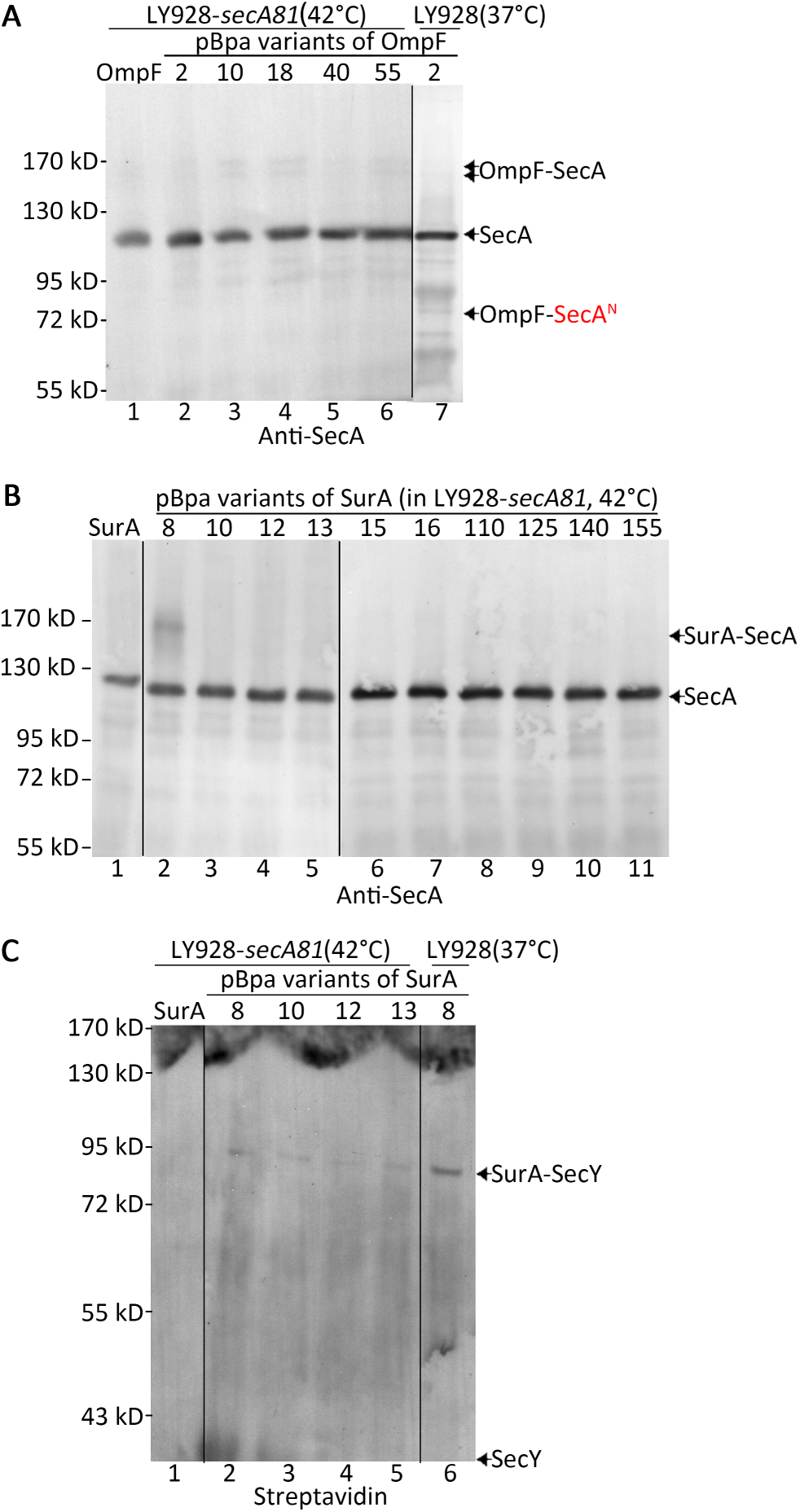
Prolonged upstream interactions of nascent β-barrel OMPs or periplasmic proteins with a defective SecA prevent their downstream interactions with SecA^N^ or SecY, respectively. (**A, B, C**) Blotting results for the detection of *in vivo* protein photo-crosslinked products of the indicated pBpa variants of nascent polypeptides of OmpF (**A**) or SurA (**B, C**) in the LY928-*secA81* mutant cells that had been cultured at the non-permissive temperature of 42°C for 1 hour, probed with antibodies against SecA (**A, B**), or with streptavidin-AP conjugate against the Avi-tag linked to SecY (**C**). As a matter of fact, samples in lane 7 of panel **A** and lane 6 of panel **C** were respectively identical to those in lane 3 of **Fig. 1B** and lane 2 of **Fig. S1B**, and were placed here to solely indicate the positions of OmpF-SecA^N^ (**A**) and SurA-SecY (**C**). Here, residue position was numbered by counting the signal peptide residues. The protein samples were resolved by SDS-PAGE before subjecting to the blotting analyses. Positions of SecA, OmpF-SecA, SurA-SecA or SurA-SecY are indicated on the right of the gels; while positions of the molecular weight markers are indicated on the left of the gels. Cells expressing the wild type OmpF or SurA (with no pBpa incorporation) were analyzed here as negative controls for the photo-crosslinking analysis (lanes 1 in **A**, **B** and **C**). The SecY protein in lanes 2-5 in panel A mobilized to the bottom edge of the gel, thus hardly visible.

Interestingly, these immunoblotting results also revealed that the photo-crosslinked products between SecA^N^ and nascent OmpF indeed became hardly detectable (lanes 1-6, **Fig. 4A**) and those between SurA and SecY likewise became significantly reduced (lanes 1-5, **Fig. 4B**) in these LY928-*secA81* cells. This was unlikely resulted from a functional defect of SecA^N^, because the assembled forms of SecA^N^ (largely produced when the cells were cultured at the permissive temperature of 37°C) were still clearly detected in these cells that had been placed at the non-permissive temperature of 42°C for only 1 h (lane 2 in **Fig. S4A**). Besides, the total amount of SecA^N^ in these LY928-*secA81* cells was still largely comparable to that detected in cells continuously cultured at 37°C also for 1 h (**Fig. S7**).

Collectively, these observations strongly suggested that, during their biogenesis, nascent polypeptides of periplasmic proteins and β-barrel OMPs would first directly but transiently interact with SecA (in the cytoplasm) before they respectively interact with SecY and SecA^N^ translocons (in the inner membrane) in wild type bacterial cells. It follows that their prolonged upstream interactions with a defective SecA (SecA81) prevented their downstream interactions with SecA^N^ or SecY in the mutant LY928-*secA81* cells that were cultured under the non-permissive temperature.

### The SecA^N^ protein directly interacts with the outer membrane-integrated β-barrel assembly machine (BAM) complex in living cells

As part of our efforts to provide further evidences to support our conclusion that SecA^N^ functions as the transmembrane protein-conducting channel for nascent β-barrel OMPs, we next sought to elucidate whether or not SecA^N^ interacts with protein factors that have been known to participate in β-barrel OMP biogenesis. In particular, we assessed whether SurA, being the major molecular chaperone that are located in the periplasmic compartment (*29-33*), and BamA, being the major component of the β-barrel assembly machine (BAM) complex which is integrated in the outer membrane but extends to the periplasm (*16-18*) interacted with SecA^N^. To achieve this goal, we first tried to find out whether pBpa individually introduced in SecA at 6 residue positions (i.e., 47, 300, 530, 863, 868 or 896), that were previously reported to be ones likely exposed to the periplasmic compartment (*65*), was able to mediate photo-crosslinking with SurA and/or BamA. Of note, here only residues 47 and 300 are present in SecA^N^, with the other four variants being also prepared and analyzed for the purpose of comparison or somehow as negative controls.

On the one hand, we failed to detect any putative photo-crosslinked product between any of these 6 pBpa variants of SecA and SurA in the LY928 cell (**Fig. S9**), suggesting that SecA^N^ on the inner membrane may not directly interact with SurA protein. On the other hand, remarkably, a putative photo-crosslinked product between SecA^N^ (but not SecA) and BamA, with a molecular mass of ~140 kD, was detected when pBpa was placed at residue position 47 in SecA (filled arrowhead, lane 1 in **Fig. 5A**). As expected, photo-crosslinked product was not detected between SecA^N^ and BamA when pBpa was placed at residue positions 530, 863, 868 or 896, none of which would be present in SecA^N^ (lanes 3-6, **Fig. 5A**). Additionally, no corsslinked product between full length SecA and BamA, a combined molecular size of them would be ~197 kD (with BamA being ~95 kD and SecA ~102 kD) was detected (**Fig. 5A**).

**Figure 5.**
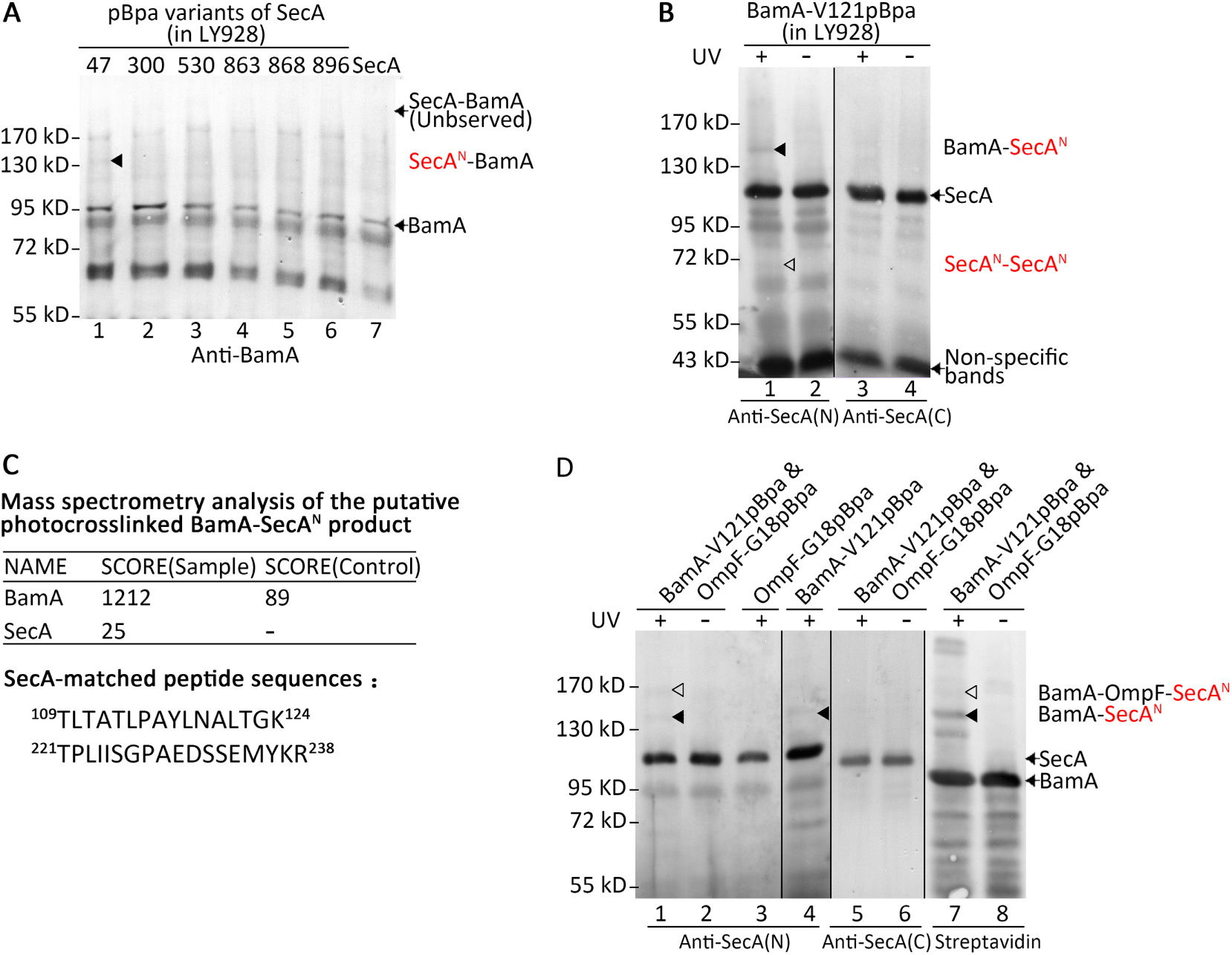
SecA^N^ directly interacts with BamA upon translocating nascent β-barrel OMPs in living cells. (**A**) Immunoblotting results for the detection of *in vivo* protein photo-crosslinked products of the indicated pBpa variants of SecA in LY928 cells, probed with antibodies against BamA. (**B**) Immunoblotting results for the detection of *in vivo* photo-crosslinked products of BamA-V121pBpa in LY928 cells, probed with antibodies against the N-terminal (lanes 1 and 2) or C-terminal region (lanes 3 and 4) of SecA. (**C**) Results of mass spectrometry analysis on the putative photo-crosslinked BamA-SecA^N^ product formed by BamA-V121pBpa. Shown are the protein scores for BamA and SecA, as well as the two matched peptide sequences from SecA. (**D**) Blotting results for the detection of photo-crosslinked products formed in the LY928 cells expressing the indicated pBpa variants of BamA and/or OmpF, probed with antibodies against the N-terminal (lanes 1-4) or C-terminal region of SecA (lanes 5 and 6), or by streptavidin-AP conjugate against the Avi-tag linked to BamA (lanes 7 and 8). For **A**, **B** and **D**, protein samples were resolved by SDS-PAGE before subjecting to the blotting analysis. Indicated on the right of the gels are positions of BamA, SecA, SecA^N^-SecA^N^, SecA^N^-BamA, BamA-SecA^N^ or BamA-OmpF-SecA^N^, on the left are positions of the molecular weight markers.

We also tried to verify this putative interaction between SecA^N^ and BamA by performing a reciprocal photo-crosslinking analysis such that the pBpa was individually placed at 9 residue positions in the periplasmic domains of BamA (*37, 38*). The immunoblotting results, displayed in **Fig. 5B**, revealed the same photo-crosslinked BamA-SecA^N^ product, again of ~ 140 kD, that could only be detected by antibodies against the N-terminal region (lane 1, filled arrowhead) but not by antibodies against the C-terminal region (lane 3) of SecA, when pBpa was placed at residue position 121 or 129, both of which are located in the POTRA 2 domain of BamA (only the results for BamA-V121pBpa is displayed in **Fig. 5B**). Interestingly, the putative photo-crosslinked SecA^N^ dimer was once again detected here, although as a faint band (lane 1, unfilled arrowhead, **Fig. 5B**)

We further isolated this putative photo-crosslinked BamA-SecA^N^ product by affinity chromatography (via the Avi-tag linked to the BamA-V121pBpa variant) and subjected it to mass spectrometry analysis. Although this indeed identified the presence of both BamA and SecA in this complex, the score for SecA was rather low and the two matched peptide fragments of SecA were both derived from the N-terminal region of SecA (**Fig. 5C**). These results were in good agreement with our expectation that it was SecA^N^, rather than SecA, that was present in the photo-crosslinked product. Taken together, these data strongly suggested that SecA^N^ directly interacted with the BamA protein in the periplasm.

Our *in vivo* photo-crosslinking data described above indicated that both OmpF (**Fig. 1B**) and BamA (**Figs. 5A** and **5B**) interacted with SecA^N^. In light of this, we then assessed whether a ternary BamA-OmpF-SecA^N^ complex is formed in living cells. For this purpose, we performed a dual *in vivo* protein photo-crosslinking analysis by co-expressing both OmpF-G18pBpa (that forms photo-crosslinked product with SecA^N^, as shown in **Fig. 1B**) and BamA-V121pBpa (that also forms photo-crosslinked product with SecA^N^, as shown in **Fig. 5B**) in LY928 cells. Blotting results, shown in **Fig. 5D**, did reveal the presence of such a photo-crosslinked ternary BamA-OmpF-SecA^N^ complex, of ~ 170 kD, which could be detected either with antibodies against the N-terminal region of SecA (lane 1, unfilled arrowhead), or with streptavidin-AP conjugate against the Avi-tag linked to BamA (lane 7, unfilled arrowhead), but not with antibodies against the C-terminal region of SecA (lane 5). As expected, the binary BamA-SecA^N^ complex, of ~ 140 kD, was meanwhile clearly detected in the sample (lanes 1, 4 and 7, **Fig. 5D**, filled arrowheads). Collectively, these results once again strongly suggested that SecA^N^ may function by interacting with BamA upon translocating nascent β-barrel OMPs across the inner membrane and then to the outer membrane.

## Discussion

This study was conducted in an initial attempt to clarify whether nascent polypeptides of both β-barrel OMPs and periplasmic proteins are translocated across the inner membrane indeed through the SecYEG translocon in living cells, as commonly believed and documented (*6, 13, 26, 48, 49*). Our *in vivo* protein photo-crosslinking analysis mediated by a genetically incorporated unnatural amino acid, however, did not reveal any direct interaction between nascent β-barrel OMPs and SecY (**Fig. 1A**), although confirmed a direct interaction between nascent periplasmic proteins and SecY (**Fig. S1A**). We then demonstrated the following. First, nascent β-barrel OMPs did, but nascent periplasmic proteins did not, directly interact with a shortened version of SecA, designated as SecA^N^, in living cells (**Figs. 1B, 1C** and **S1B**). Second, SecA^N^ was specifically present in the membrane fraction and apparently existed as homo-oligomers (**Fig. 2**). Third, SecA^N^ contains a putative transmembrane domain that contains a GXXXG motif known for mediating protein-protein interactions in biomembranes (**Table 1**). Fourth, the processing of precursors of β-barrel OMPs was severely affected in cells producing assembly-defective SecA^N^ variants due to mutations introduced in the GXXXG motif (**Fig. 3** and **Table 2**). Fifth, prolonged interactions of nascent β-barrel OMPs or periplasmic proteins with a defective SecA (SecA81) prevented their downstream interactions with SecA^N^ or SecY, respectively (**Fig. 4**). Sixth, the SecA^N^ protein directly interacted with the outer membrane-integrated β-barrel assembly machine (BAM) complex in living cells (**Fig. 5**). Seventh, precursor processing of β-barrel OMPs relied largely on ATP but hardly on the transmembrane proton gradient, while that of periplasmic proteins relied on both as energy sources (**Fig. S10**).

Our new findings, in combination with previous revelations (*29-36, 66-68*), strongly imply that SecA^N^ and SecYEG are responsible for translocating nascent polypeptides of β-barrel OMPs and periplasmic proteins, respectively, across the inner membrane. Here, as schematically illustrated in **Fig. 6**, we emphasize the following points. First, as the translocon of nascent β-barrel OMPs, SecA^N^, likely exists as a homo-oligomer (arbitrarily shown as a dimer in **Fig. 6**, although its exact oligomeric form remains to be elucidated). Second, the full length form of SecA in the cytoplasm (being membrane associated or free in the cytoplasm) is assumed to deliver the signal peptide-containing nascent β-barrel OMPs and nascent periplasmic proteins to the membrane-integrated translocons SecA^N^ and SecYEG, respectively. Third, SecA^N^ is proposed to be part of the supercomplex that we revealed earlier as one being responsible for the biogenesis of β-barrel OMPs in living cells and spanning the cytoplasm, the inner membrane, the periplasm and the outer membrane (*33*).

**Figure 6.**
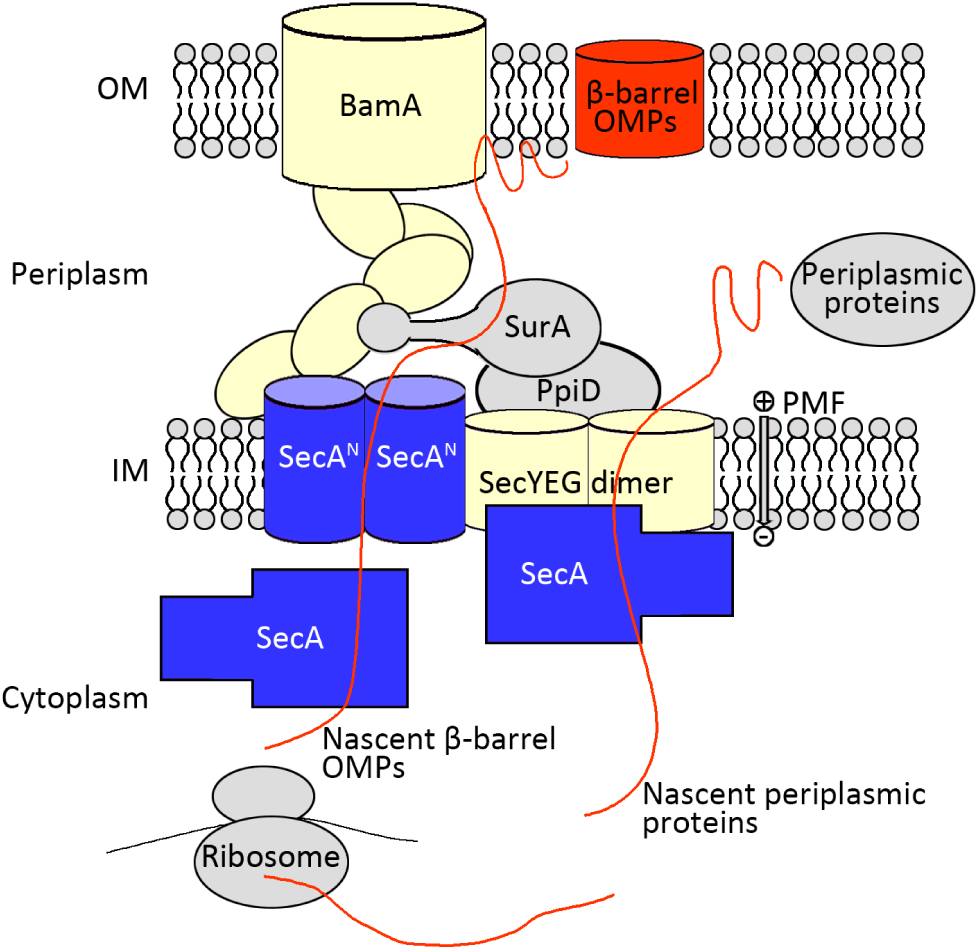
A schematic illustration on the translocation of nascent polypeptides of β-barrel OMPs and periplasmic proteins across the inner membrane in Gram-negative bacteria by using SecA^N^ and SecYEG as translocons, respectively.

As regards to how SecA^N^ monomers, if initially being produced as a soluble form, are converted to the membrane-integrated oligomers, the case of pore-forming toxins might be referred to. It has been long recognized that these pore-forming toxins, initially produced as soluble and inactive forms by many pathogenic bacteria, are able to assemble in cell membranes to form large multimeric pores that will allow protein molecules to be transported through (*69*). It remains to be elucidated whether SecA^N^ shares similar properties with these toxins. We noticed that part of the GXXXG motif is actually located in a β-strand structure in the full length form of SecA, as shown in **Fig. S5**. Whether there exists a β-strand to α-helix conformational conversion similar to what occurs in the prion proteins (*69*) is also worth further exploration.

One may ask why the SecA^N^ protein has not been unveiled over the past decades by people working on SecA or on bacterial protein translocations. The following points may explain this. First, SecA^N^ might have been considered as a degradation product of SecA. Second, SecA^N^, almost wholly resides in the membranes and presents in the whole cell extract at a level far lower than that of SecA, thus might have escaped any attention. Third, since SecA^N^ is tightly integrated into the inner membrane, it might have thus escaped the detection by immunoprecipitation analysis as commonly performed in the early days (*70*). Fourth, in using the antibodies that were made widely available to the community (*71*), we noticed the existence of multiple non-specific protein bands at the position around SecA^N^, which might have added further difficulties for it to be noticed (as shown in **Fig. 1B**).

Despite of these, we nevertheless searched the literatures seeking for early hints on the existence of SecA^N^ in bacterial cells. Interestingly, we did notice the report of a SecA form that had a monomeric size very close to SecA^N^ (~ 45 kD) and could be immunoprecipitated from the lysates of *E. coli* cells by antibodies against the N-terminal region of SecA (as shown in lanes 7 and 8 of **Fig. 2** in (*72*)) or be detected in the inner membrane fraction by antibodies against the full length SecA (**Fig. 4** in (*65*)). Moreover, it was reported that the N-terminal fragments of about 240 residues of the SecA protein (which would contain the transmembrane domain we identified here) from either *E coli* (*57*) or *B. Subtilis* (*73*) could be integrated into the membrane. Many issues remain to be resolved on the structure and function of SecA^N^. For example, how SecA^N^, apparently existing as homo-oligomers, forms the protein conducting channel in the inner membrane? Is SecA^N^ directly generated during the translation process or derived from the full length form of SecA by protease cleavage? Does the protein SecM, which has been known to regulate the translation of SecA in cells (*64*), play any role in the production of SecA^N^? How SecA^N^ on the inner membrane interacts and works with the full length SecA present in the cytoplasm?

Our new findings reported here mad it necessary to reappraise the long-held view that SecYEG is directly responsible for translocating nascent polypeptides of both β-barrel OMPs and periplasmic proteins in bacterial cells (*6, 26, 48, 49*). In retrospect, this perception has been formed mainly from results of genetic and *in vitro* reconstitution studies (*13, 74*), as we described above. Consistent with what was reported before, our data also indicated that SecY, though not serving as the actual conducting channel, does play certain role in translocating nascent β-barrel OMPs across the inner membrane in living cells (**Fig. S2**). This might be explained as such that SecYEG is responsible for translocating such periplasm-located quality control factors as SurA and DegP (*29-31, 33, 47*), which play important roles for escorting nascent β-barrel OMP across the periplasm.

Unresolved issues regarding the relationship between SecYEG and SecA^N^ include the following. Do SecA^N^ and SecYEG have any functional coordination in respectively translocating β-barrel OMPs and periplasmic proteins, such that, for example, allowing the latter two groups of proteins to be produced in a certain coordinated manner? How does the cytoplasmic full length form of SecA effectively partition nascent β-barrel OMPs and nascent periplasmic proteins to SecA^N^ and SecYEG, respectively? The molecular mechanism for SecA^N^ to function as a protein transcolon in the inner membrane, as well as the structural and functional interaction between SecA^N^ and SecYEG translocons certainly merit future explorations.

## Methods

### Bacteria strains and plasmid construction

All bacteria strains and plasmids used in this research are respectively listed in **Tables S1** and **S2**. The pYLC-OmpF, pYLC-BamA, pYLC-SurA or pYLC-SecA plasmid was constructed by isolating the gene (including its promoter) fragment via PCR, using the *E. coli* genomic DNA as template; the DNA fragment was inserted into the pYLC plasmid vector through restriction enzyme-free cloning (*75*). The pYLC plasmid is a low-copy plasmid that we derived from the pDOC plasmid (*33*). The site-specific replacement of a certain codon by the TAG amber codon in a particular gene (to introduce a pBpa residue at the position) was performed using the phusion site-directed mutagenesis kit (New England Biolabs, Massachusetts, USA).

### The *in vivo* protein photo-crosslinking mediated by pBpa

The pYLC-OmpF or pYLC-SurA plasmid carrying an introduced TAG codon in the *ompF* or *surA* gene was respectively expressed in LY928-∆*ompF* or LY928-∆*surA* cells. The LY928 strain was generated by us for pBpa incorporation purposes, as described before (*33*). The cells were cultured at 37°C in LB medium containing pBpa (200 μM) to the mid-log phase (with an OD_600_ of ~0.8-1.0) before irradiated with UV light (365 nm) for 10 min in a Hoefer UVC-500 crosslinker. The UV-irradiated cells were then harvested by centrifugation, resuspended in SDS-PAGE loading buffer before boiled for 5 min. The protein samples were subsequently resolved by SDS-PAGE before being subjected to blotting analysis.

### Separation of soluble and membrane fractions of lysed bacterial cells and UV irradiation of the separated fractions

The separation of soluble and membrane fractions of lysed bacterial cells was performed according to methods described before (*56, 57*) with some modifications. Briefly, the LY928 cell was cultured to the mid-log phase (with an OD_600_ of ~0.8-1.0), collected by centrifugation, resuspended and washed with PBS buffer, lysed (in PBS buffer containing 0.1 mM PMSF) by sonication before centrifuged at 10,000 x g for 5 min to remove the cell debris and unlysed cells. The supernatant was then centrifuged at 22,000 x g for 20 min to separate the soluble and membrane fractions. Both fractions were irradiated with UV light (365 nm) for 10 min in a Hoefer UVC-500 crosslinker, before being subjected to SDS-PAGE and blotting analyses.

### Mass spectrometry analysis

Photo-crosslinked product of BamA-V121pBpa was purified by affinity chromatography using streptavidin resin. The eluted sample was resolved by SDS-PAGE before the gel was subjected to either blotting analysis or Coomassie Blue staining. The protein bands around 140 kD was then excised from the gel and applied for liquid chromatography-tandem mass spectrometry (LC-MS/MS) identification as we described before (*33*).

### Chloramphenicol treatment of the LY928-*secA81* cells transformed with the plasmid expressing the Avi-SecA-G155F variant

A single colony of LY928-*secA81* cells transformed with the plasmid expressing Avi-SecA-G155F variant was inoculated in 3 mL of LB medium containing 50 μg/mL Kanamycin, cultured at the permissive temperature of 37°C to an OD_600_ of ~ 0.3-0.4, shifted to and incubated at the non-permissive temperature of 42°C for 3 hour before treated with chloramphenicol (34 μg/mL) for 30 min or 1 hour. Treated cells were then collected by centrifugation, resuspended in loading buffer, incubated at 37°C for 10 min before applied to semi-native SDS-PAGE analysis. Protein samples were resolved by SDS-PAGE before being subjected to blotting analysis.

## Acknowledgements

We thank the Keio Collections for providing us the wild type and mutant (single-gene knockout) *E. coli* strains. We thank Professor Koreaki Ito from Kyoto Sangyo University for providing us the *secY39* mutant strain. We thank Prof. Phang-Cheng Tai (Georgia State University), Prof Senfang Sui (Tsinghua University) Prof. Xinmiao Fu (Fujian Normal University) for valuable discussions. We thank Dr. Wen Zhou at the Mass spectrometry Facility of the National Center for Protein Sciences at Peking University for assistance in performing mass spectrometry analysis. We thank Mr. Yang Liu and Dr. Jiayu Yu (members of our lab) for providing the LY928 strain and the pYLC plasmid vector. We thank Dr. Yan Wang and Dr. Rui Wang for performing mass spectrometry analysis of the photo-crosslinked products of OmpF, Ms. Jimei Zhao for constructing the plasmids expressing the pBpa variants of SurA, Ms. Pan Zou for providing us the plasmid expressing SecY-Avi (all former members of our lab). This study was supported by research grants from the National Natural Science Foundation of China (No. 31670775 and No. 31470766 to ZYC) and the national Basic Research Program of China (973 Program) (No. 2012CB917300 to ZYC as the chief scientist).

## Author Contributions

Feng Jin designed and performed the experiments. Zengyi Chang supervised this whole study. Feng Jin and Zengyi Chang analyzed the data and prepared the manuscript.

